# Direct comparison of SARS-CoV-2 variant specific neutralizing antibodies in human and hamster sera

**DOI:** 10.1101/2023.12.19.572347

**Authors:** Annika Rössler, Antonia Netzl, Ludwig Knabl, Samuel H. Wilks, Barbara Mühlemann, Sina Türeli, Anna Mykytyn, Dorothee von Laer, Bart L. Haagmans, Derek J. Smith, Janine Kimpel

## Abstract

Antigenic characterization of newly emerging SARS-CoV-2 variants is important to assess their immune escape and judge the need for future vaccine updates. As exposure histories for human sera become more and more complex, animal sera may provide an alternative for antigenic characterization of new variants. To bridge data obtained from animal sera with human sera, we here analyzed neutralizing antibody titers in human and hamster first infection sera in a highly controlled setting using the same live-virus neutralization assay performed in one laboratory. Using a Bayesian framework, we found that titer fold changes in hamster sera corresponded well to human sera and that hamster sera generally exhibited higher reactivity. Our results indicate that sera from infected hamsters are a good surrogate for the antigenic characterization of new variants.

## Introduction

Antigenic characterization is critical for tracking the evolution of severe acute respiratory syndrome coronavirus type 2 (SARS-CoV-2), assessing the immune escape of emerging SARS-CoV-2 variants, and judging the need for vaccine updates (1). This requires the measurement of neutralizing antibody titers in cohorts representing the current population immunity to validate the need for a vaccine update. Additionally, first exposure sera are essential to assess the antigenic properties of the variant in the context of previously circulating variants, without confounding by immunity from previous infections. Multi-exposure sera are unsuitable for that purpose as multiple exposures generally increase cross-neutralization and thereby obscure the underlying antigenic relationships among variants (2, 3). Antigenic characterization of virus variants can be performed using human or animal sera. By now, first exposure human sera against newly emerging SARS-CoV-2 variants are increasingly difficult to collect. For influenza virus, ferrets are the standard organism for characterizing antigenic relationships (4). Hence, bridging of human and animal data is an important task for future research and public health management of SARS-CoV-2.

## Results

Here, we directly compared neutralizing antibody titers from first exposure human and hamster sera. We selected four serum groups for this study: non-vaccinated individuals infected with the ancestral virus (614D/G, n_human_=10, n_hamster_=4), Delta (n_human_=7, n_hamster_=4), BA.1 Omicron (n_human_=17, n_hamster_=4), or BA.5 Omicron variant (n_human_=3, n_hamster_=3) (Supplementary Table 1). To minimize bias by different assays and interlaboratory variation, we analyzed neutralizing antibody titers for all samples in the same laboratory using the same focus reduction neutralization assay for 6 pre-Omicron (D614G, Alpha, Alpha-E484K, Beta, Gamma, Delta) and 4 Omicron (BA.1, BA.2, BA.5.3.2, XBB.1.5.1) variants (Supplementary Table 2). The variants and sera used had previously been published in independent studies in two different labs using different assays (2, 5, 6).

We found that while overall fold change trends for human and hamster titers were similar, variation between subjects in the same serum group was lower for hamster samples and hamster titers were generally higher (Figure 1, row 1). This was true for all serum groups. The higher variation in human samples might be explained by the natural variation between samples in this group in terms of genetic background, age, time since exposure, infecting dose, infecting virus etc. To control for this within and between serum magnitude difference, we employed a Bayesian framework which was recently used to compare SARS-CoV-2 neutralization data across different laboratories and species (7). In this framework, variations from the overall geometric mean titer for a variant in a serum group are attributed to different individual serum reactivities, reactivity differences due to species, and noise. Using this statistical framework on our experimental data allowed us to quantify serum and organism specific effects and obtain estimates for titers that fall below an assay’s limit of detection (LOD) (Figure 1, rows 2-4).

**Figure 1.**
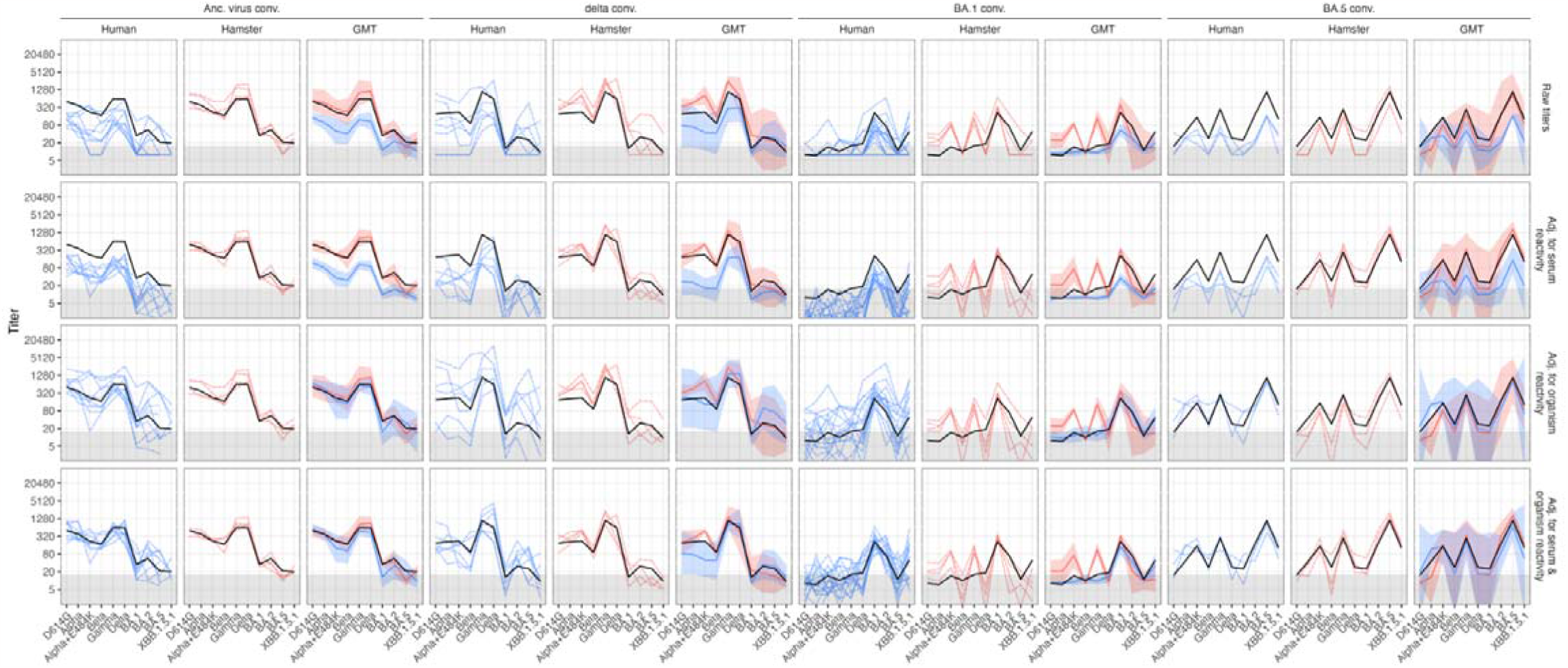
Comparison of neutralization titers in hamster and human single variant exposure sera. Human and hamster single variant exposure sera (human n=10 ancestral, n=7 delta, n=17 BA.1, n=3 BA.5; hamster n= 4 ancestral, n=4 delta, n=4 BA.1, n=3 BA.5) were analyzed for neutralizing antibodies against D614G, alpha, alpha+E484K, beta, gamma, delta, BA.1, BA.2, BA.5, XBB.1.5.1 variants using a focus reduction assay and live virus variants. To control for titer variation due to different reactivities of individual sera and estimate species-specific effects, titers were estimated using a Bayesian framework (7). The columns show human titers (blue), hamster titers (pink) and the GMT (geometric mean titers) ± +95 % CI (confidence interval) as bold colored line and shaded area for each convalescent (conv.) serum group. The black line represents the estimated Geometric Mean Titer per serum group across organisms after adjusting for serum and organism effects. The rows show from top to bottom: Raw titers with titers <LOD (limit of detection ≤ 16 indicated by grey area) set to 8 (LOD/2), titers adjusted for individual serum reactivity variation, titers adjusted for organism reactivity differences, and titers adjusted for both individual serum and organism reactivities.

Adjusting for serum reactivity effects reduced variation among individual samples as expected and showed that although titer magnitude may differ between individuals, the fold changes (and consequently the relative titer differences between variants) were very similar and independent of titer magnitude (Figure 1, row 2). We further found a systematic difference in titer magnitude between human and hamsters, and calculated hamster titers to be an average estimate of 4.85-fold higher than human titers (Figure 1, row 2, magnitude adjusted in row 3). The human-hamster titer magnitude difference estimate of 3.1-fold higher hamster titers by Mühlemann *et al*. (7) fell within the distribution’s high-density interval (Supplementary Figure 1). After controlling for both within- and between-species titer magnitude effects, human and hamster titer fold change trends were highly similar in the Ancestral virus convalescent (conv.) and BA.5 conv. serum groups, apart from lower titers against the Alpha variant in BA.5 conv. hamsters (Figure 1, row 4). We further found that hamster BA.1 sera had higher titers against Alpha+E484K and Gamma than human sera. In Delta sera, too, hamsters exhibited higher titers against Alpha+E484K and Alpha than humans, but fold change trends were otherwise similar. To summarize, fold change patterns are remarkably consistent across humans and hamsters except for the few differences described above.

## Discussion

Mühlemann *et al*. introduced a systematic framework to compare SARS-CoV-2 antigenic data from different laboratories, generated without sharing of sera or variants and using different assays (7). Here, we compared data obtained in a highly controlled setting: Human and hamster sera were measured in a single laboratory in the same assay and with the same virus stock. Mühlemann *et al*.’s framework recommends evaluation of antigenic data on four levels: Titer magnitude, titer fold changes, immunodominance patterns, and antigenic cartography (7). Although the controlled nature of our study comes with the limitation that we did not have serum groups available from both species with known immunodominance changes, such as Beta convalescent sera (7, 8), and lacked single-exposure sera from diverse variants to construct well-triangulated antigenic maps (Supplementary Figure 2), it allowed for exact comparison of titer magnitude and fold change trends. Our data indicate that titer fold changes in hamster sera correspond well to human sera, and that titers correspond well after adjusting the data for the higher reactivity generally seen in the hamster. The higher reactivity in hamster sera will consequently not influence the antigenic relationships determined by the assay, but rather provide benefit as a greater range of neutralization can be measured before values fall below an assay’s limit of detection. In the context of antigenic cartography, this greater detection range can be especially useful as it permits longer range triangulation and consequently increases map resolution (4, 9). In summary, for the sera raised, variants tested, and assay used here, sera from infected hamsters are a good surrogate for human sera for the antigenic characterization of SARS-CoV-2 variants and overcome the limitation of collecting human single variant exposure sera for newly emerging virus variants. It will be important however to continue to test the suitability of hamster sera as a model for single-exposure human sera against newer variants.

## Materials and Methods

### Experimental design

The objective of this study was to compare SARS-CoV-2 variant specific neutralizing antibody titers in human and hamster samples. For this purpose, serum samples were collected from non-vaccinated humans or hamsters recovered from a single infection with ancestral (humans: 614D, hamsters: D614G), Delta, BA.1 Omicron, or BA.5 Omicron variant. Titers of neutralizing antibodies against SARS-CoV-2 variants were determined in human and hamster sera in the same neutralization assay in a single laboratory. To control for titer variation due to different reactivities of individual sera and estimate species-specific effects a Bayesian framework was employed (7).

### Human samples

Sera were collected from 37 individuals after SARS-CoV-2 single variant exposure. In more detail, we analyzed the samples from individuals after infection with ancestral (n=10), Delta (n=7) or Omicron BA.1 (n=17) or BA.5 (n=3) variant and study cohorts are characterized in Supplementary Table 1. The ethics committee (EC) of the Medical University of Innsbruck has approved sample collection with EC numbers: 1100/2020, 1111/2020, 1330/2020, 1064/2021, 1093/2021, 1168/2021, 1191/2021, 1197/2021, and 1059/2022. Informed consent has been obtained from study participants.

### Animal samples

Female Syrian golden hamster sera was obtained as described previously (6, 10). Briefly, animals were inoculated intranasally with 1 × 10^5^ plaque-forming units of ancestral (n=4) or Omicron BA.5 (n=3), or 5 × 10^4^ of Delta (n=4) or Omicron BA.1 (n=4). Omicron BA.5 animals were euthanized at 21 days post infection, whereas all other animals were euthanized at 26 days post infection, at which point serum was collected. This research was in compliance with the Dutch legislation for the protection of animals used for scientific purposes (2014, implementing EU Directive 2010/63). This research was conducted either at Erasmus MC (approved OLAW Assurance no. A5051-01, study protocol no. 17-4312 approved by institutional Animal Welfare Body) or at Viroclinics Biosciences B.V., Viroclinics Xplore (license number AVD27700202114492-WP35).

### Neutralization assay

Human and hamster samples were tested for neutralization against a panel of 10 live SARS-CoV-2 isolates, which included several pre-Omicron variants (D614G, Alpha, Alpha with additional E484K mutation, Beta, Gamma and Delta), three Omicron variants (BA.1, BA.2 and BA.5) as well as a recombinant lineage (XBB.1.5.1). Details on used virus isolates are shown in Supplementary Table 2. To analyze neutralization titer, we performed a focus forming assay as previously described (2). Therefore, four-fold dilutions of heat-inactivated sera were incubated with SARS-CoV-2 isolates for 1 h at 37°C and subsequently transferred to Vero-cells overexpressing TMPRSS2 and ACE2. The virus/sera mix was replaced by fresh medium 2 h after infection and cells were fixed with absolute ethanol further 8h later. Infected cells were visualized by immunofluorescence staining. Results of the human samples were developed using a SARS-CoV-2 convalescent plasma as primary and a goat anti-human IgG Alexa Fluor 488 secondary antibody (ThermoFisher Scientific #A48276) (2). For hamster samples a SARS-CoV-2 Nucleocapsid antibody (SinoBiological #40588) followed by a goat anti-rabbit IgG Alexa Fluor 488 antibody (ThermoFisher Scientific #A32731) was used (6). Infected cells were counted using an immunospot reader and continuous neutralization titers (IC_50_) were calculated by non-linear regression (GraphPad Prism Software 9.0.1, Inc., La Jolla, CA, USA). Titers ≥16,384 were set to 16,384. Neutralization titers ≥16 were considered positive and negative titers were set to half the detection limit, i.e. 8. For Bayesian modelling and antigenic cartography (Supplementary Figure 2), titers <16 were set to “<16”.

### Titer adjustments

A recent study comparing SARS-CoV-2 neutralization data from different assays, species and laboratories found that reactivity patterns were similar, but titer magnitude varied by species (7). The study employed a modelling approach to control for magnitude differences, which was adapted here: Each titer is modelled as a combination of geometric mean titer per variant and serum group, a reactivity effect of each individual serum explained by individual high or low immune responses, and a species-specific magnitude difference.

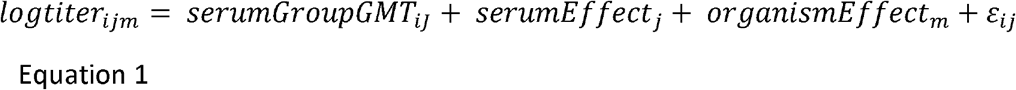

In Equation 1, the *serumGroupGMT* refers to the log2 titer of antigen *i* in serum group *J*, the serumEffect corresponds to the reactivity bias of serum *j*, the organismEffect to the reactivity bias of organism *m*, and log2 normally distributed noise for each measurement, where the standard deviation is assumed to differ between organisms.

The model was based on the cmdstanr model used by Mühlemann *et al*. (7) (R version 4.2.2 (11), cmdstanr version 0.5.3 (12)), and priors for the standard deviation parameters were chosen based on their values (inverse gamma distribution, shape = 3, scale = 1.5). For modelling both serum and organism reactivity effects, the following distributions were used: serumGroupGMT: N(3, 20), serumEffect: N(0, 10), organismEffect: N(0, 2). For modelling only serum or organism effects (Supplementary figures 1-3), the standard deviation of the excluded effect was set to 1e-3 to penalize any deviation from the mean 0. The mean of the posterior distributions when modelling both organism and serum reactivity was used to adjust the raw titers (Supplementary Figure 1). cmdstanr’s sampling, rather than its optimization function, was used as a comparison of values revealed a small difference between posterior means and optimized values (Supplementary Figures 1, 3-4). This may happen when the posterior distribution is not convex and the optimization algorithm gets stuck in a local optimum instead of a global one.

All code can be found in the manuscript’s GitHub repository (13): https://github.com/acorg/roessler_netzl_et_al2023a.git

## Acknowledgments

We thank Albert Falch and Verena Pittl for their excellent technical support.

## Funding

NIH NIAID Centers of Excellence for Influenza Research and Response (CEIRR) contract 75N93021C00014 as part of the SAVE program (JK, DJS, BLH, AN, SHW, and ST)

European Union’s Horizon 2020 research and innovation program under grant agreement No. 101016174 (JK)

Austrian Science Fund (FWF) with the project number P35159-B (JK)

Gates Cambridge Trust (AN)

## Author contributions

AR and AN contributed equally to this work. AR designed and performed neutralization assays. AN performed the antigenic cartography and Bayesian Modeling, which was supervised by DJS. LK, AM, BLH collected and contributed samples. SHW, BM, ST provided methodology to analyze data. DvL co-designed the study. JK designed and supervised the study and wrote the manuscript together with AN. All authors edited and reviewed the manuscript prior to submission.

## Data and materials availability

All data and code is publicly available in the manuscript’s GitHub repository (13) (https://github.com/acorg/roessler_netzl_et_al2023a.git).

## References

1. DeGrace MM, Ghedin E, Frieman MB, Krammer F, Grifoni A, Alisoltani A, et al. Defining the risk of SARS-CoV-2 variants on immune protection. Nature. 2022;605(7911):640–52.

2. Rössler A, Netzl A, Knabl L, Schäfer H, Wilks SH, Bante D, et al. BA.2 and BA.5 omicron differ immunologically from both BA.1 omicron and pre-omicron variants. Nat Commun. 2022;13(1):7701.

3. Netzl A, Tureli S, LeGresley E, Mühlemann B, Wilks SH, Smith DJ. Analysis of SARS-CoV-2 Omicron Neutralization Data up to 2021-12-22. bioRxiv. 2022:2021.12.31.474032.

4. Smith DJ, Lapedes AS, de Jong JC, Bestebroer TM, Rimmelzwaan GF, Osterhaus AD, et al. Mapping the antigenic and genetic evolution of influenza virus. Science. 2004;305(5682):371–6.

5. Mykytyn AZ, Rissmann M, Kok A, Rosu ME, Schipper D, Breugem TI, et al. Antigenic cartography of SARS-CoV-2 reveals that Omicron BA.1 and BA.2 are antigenically distinct. Sci Immunol. 2022;7(75):eabq4450.

6. Mykytyn AZ, Rosu ME, Kok A, Rissmann M, van Amerongen G, Geurtsvankessel C, et al. Antigenic mapping of emerging SARS-CoV-2 omicron variants BM.1.1.1, BQ.1.1, and XBB.1. Lancet Microbe. 2023;4(5):e294–e5.

7. Mühlemann B, Wilks SH, Baracco L, Bekliz M, Carreño JM, Corman VM, et al. Comparative Analysis of SARS-CoV-2 Antigenicity across Assays and in Human and Animal Model Sera. bioRxiv. 2023:2023.09.27.559689.

8. Wilks SH, Mühlemann B, Shen X, Türeli S, LeGresley EB, Netzl A, et al. Mapping SARS-CoV-2 antigenic relationships and serological responses. Science. 2023;382(6666):eadj0070.

9. Rössler A, Netzl A, Knabl L, Bante D, Wilks SH, Borena W, et al. Characterizing SARS-CoV-2 neutralization profiles after bivalent boosting using antigenic cartography. Nat Commun. 2023;14(1):5224.

10. Mykytyn AZ, Rissmann M, Kok A, Rosu ME, Schipper D, Breugem TI, et al. Antigenic cartography of SARS-CoV-2 reveals that Omicron BA.1 and BA.2 are antigenically distinct. Science Immunology. 2022;7(75):eabq4450.

11. R Core Team. R: A Language and Environment for Statistical Computing Vienna, Austria: R Foundation for Statistical Computing; 2022 [Available from: https://www.R-project.org/.

12. Gabry J, Češnovar R, Johnson A. cmdstanr: R Interface to ‘CmdStan’ 2023 [Available from: https://mc-stan.org/cmdstanr/, https://discourse.mc-stan.org.

13. Netzl A. acorg/roessler_netzl_et_al2023a: Initial submission of manuscript (v.1.0) 2023 [Available from: 10.5281/zenodo.10003230.

