## Supplementary Materials for "Direct comparison of SARS-CoV-2 variant specific neutralizing antibodies in human and hamster sera"

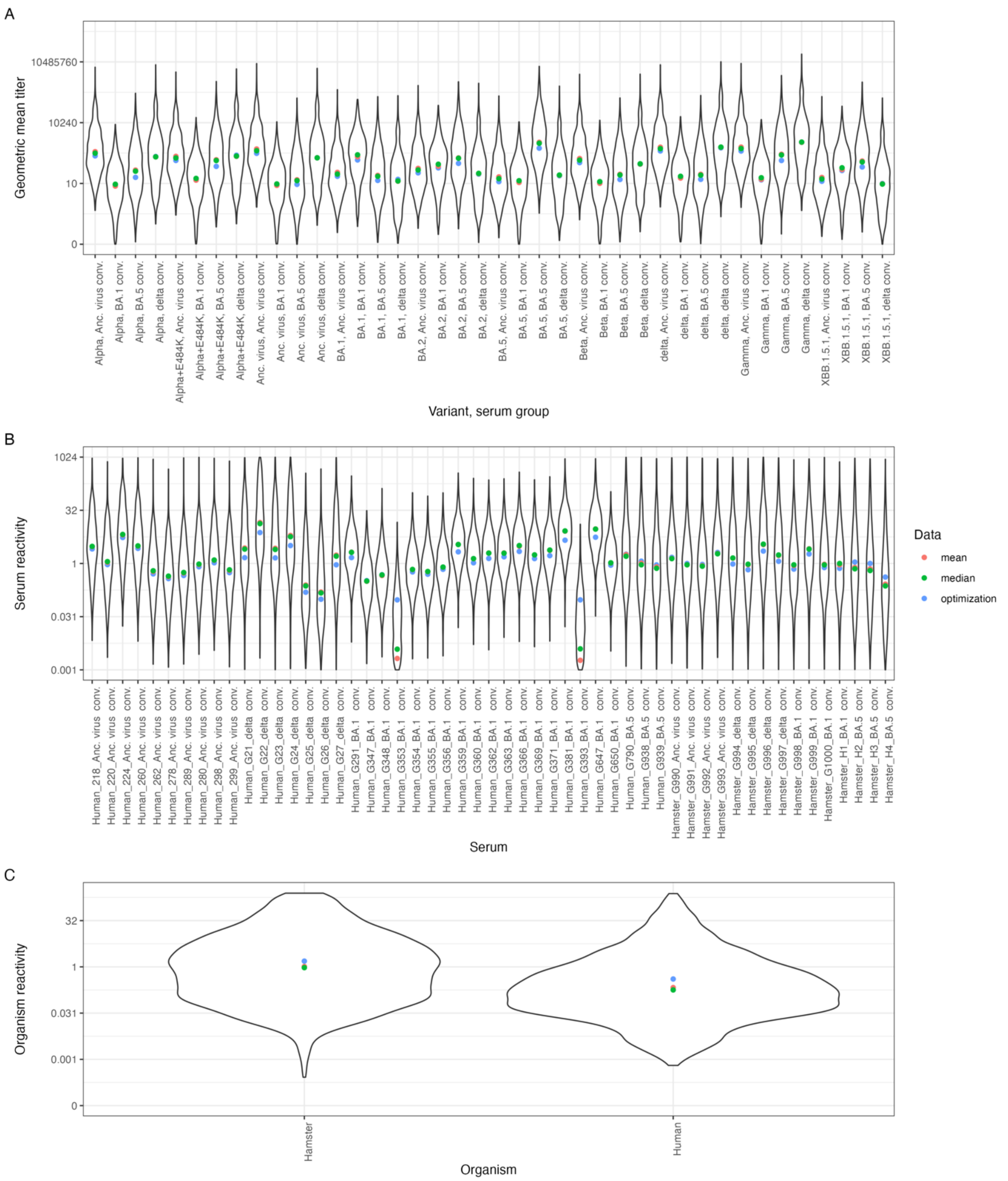


### Fig. S1.

**Posterior distribution when modelling serum and organism reactivity.** Posterior distributions of draws for **A)** variant geometric mean titers, **B)** serum reactivities, and **C)** organism reactivities are shown. The mean of the distribution is shown in pink, the median in green and the optimized value in blue. The reactivity corresponds to a fold change of titers against all antigen variants.

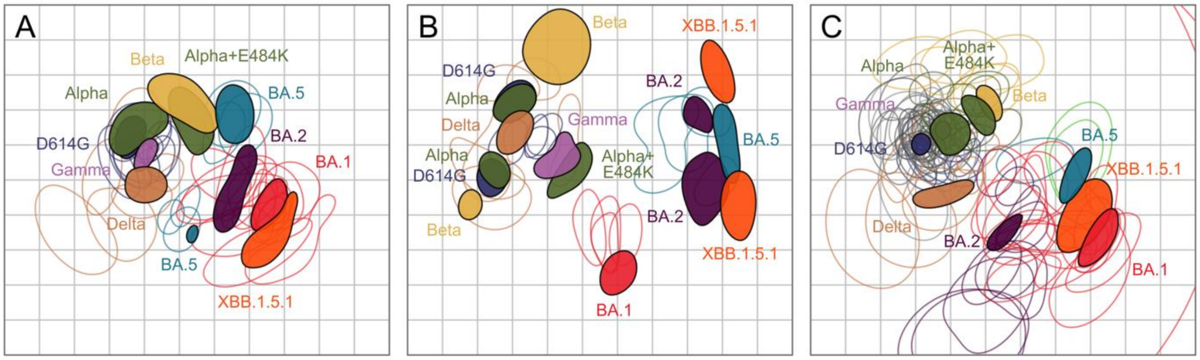


**Fig. S2.**

**Positional uncertainty in human and hamster antigenic maps. A)** Human and **B)** hamster antigenic maps were constructed in Racmacs (8) (v 1.1.35) with 1000 optimizations in 2 dimensions. **C)** Human map with additional serum groups (two dose Ancestral variant vaccinated, Beta convalescent, Alpha convalescent, BA.2 convalescent) subset to variants in this comparison (originally published by Rössler *et al*. (9). Positional variation of human and hamster sera was determined by Bayesian bootstrapping, where sera and antigen measurements are weighted by random draws from a Dirichlet distribution, such that individual sera and antigens contribute to the map optimization with different weights. 500 bootstrap repeats were performed with 1000 optimizations each. The colored regions mark 68 % (one standard deviation) of the positional variation for each variant (filled shapes) and sera (open shapes). Multiple shapes per variant, as seen in the human map for BA.5 and in the hamster map for D614G, Alpha, BA.2 and XBB.1.5.1, indicate that multiple optima exists and that the map is not well triangulated. Additional serum groups increase map triangulation and positional certainty as shown in **C.**

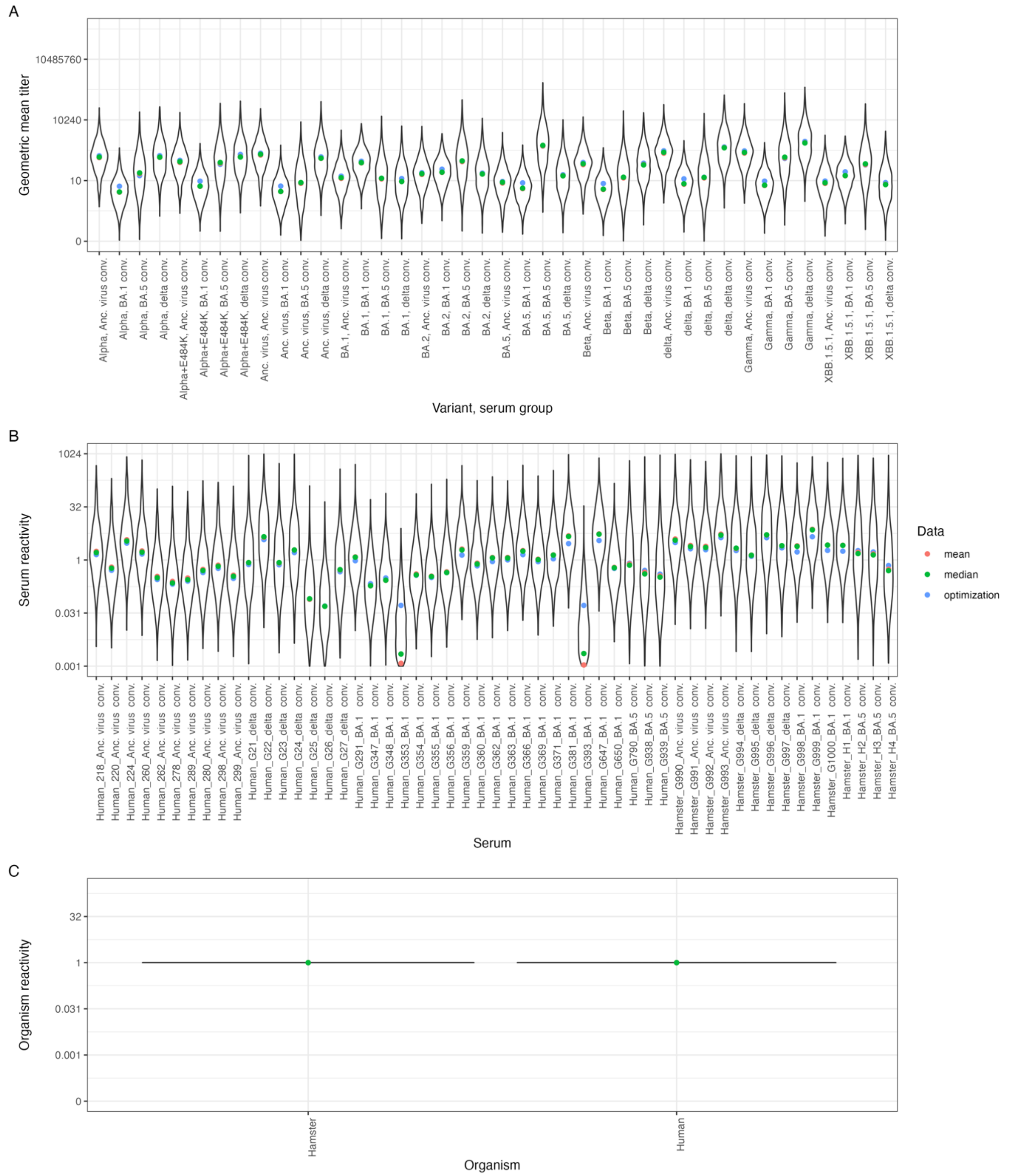


**Fig. S3.**

**Posterior distribution when modelling only serum reactivity.** Posterior distributions of draws for **A)** variant geometric mean titers, **B)** serum reactivities, and **C)** organism reactivities are shown. The mean of the distribution is shown in pink, the median in green and the optimized value in blue. The reactivity corresponds to a fold change of titers against all antigen variants.

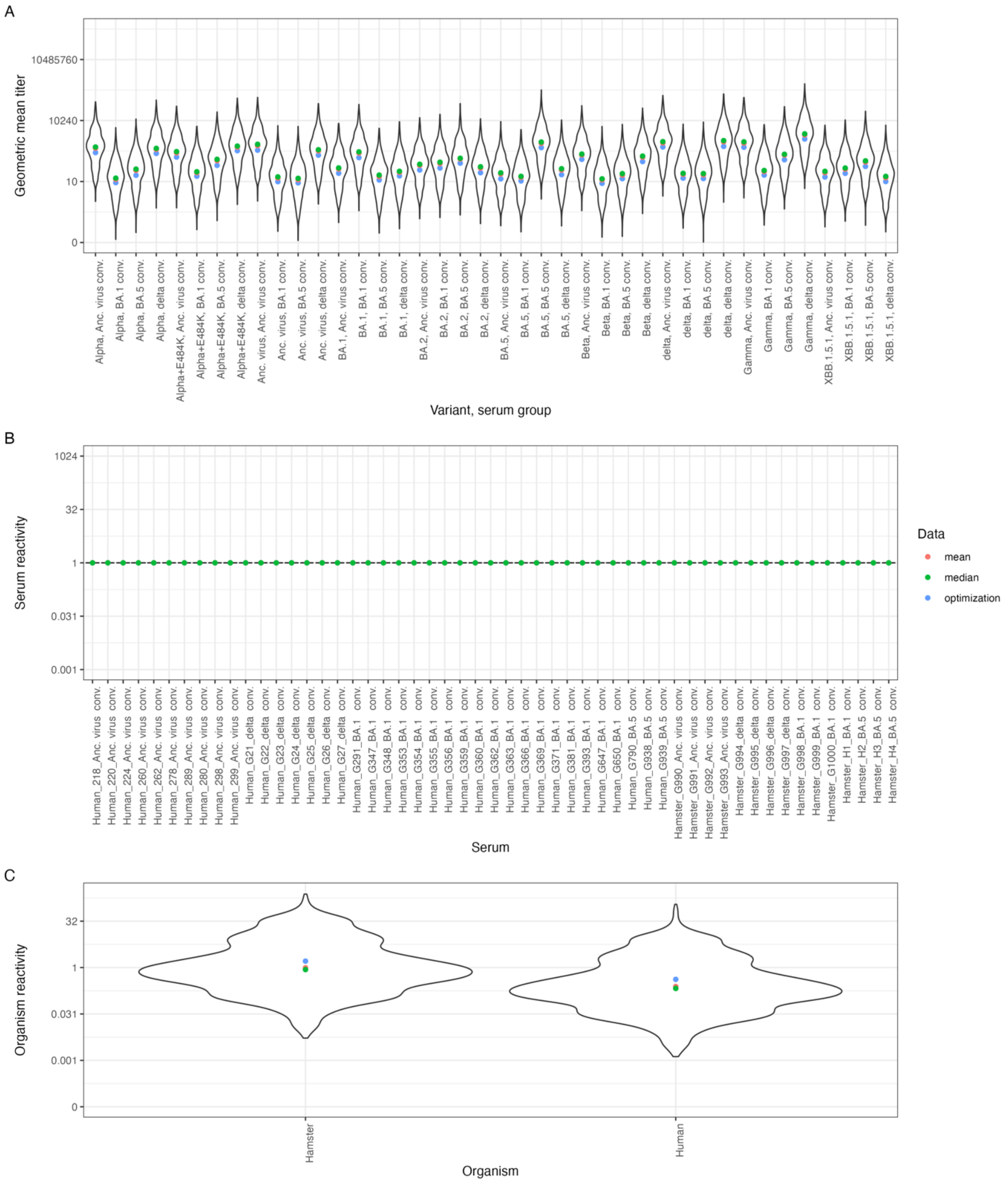


**Fig. S4.**

**Posterior distribution when modelling only organism reactivity.** Posterior distributions of draws for **A)** variant geometric mean titers, **B)** serum reactivities, and **C)** organism reactivities are shown. The mean of the distribution is shown in pink, the median in green and the optimized value in blue. The reactivity corresponds to a fold change of titers against all antigen variants.

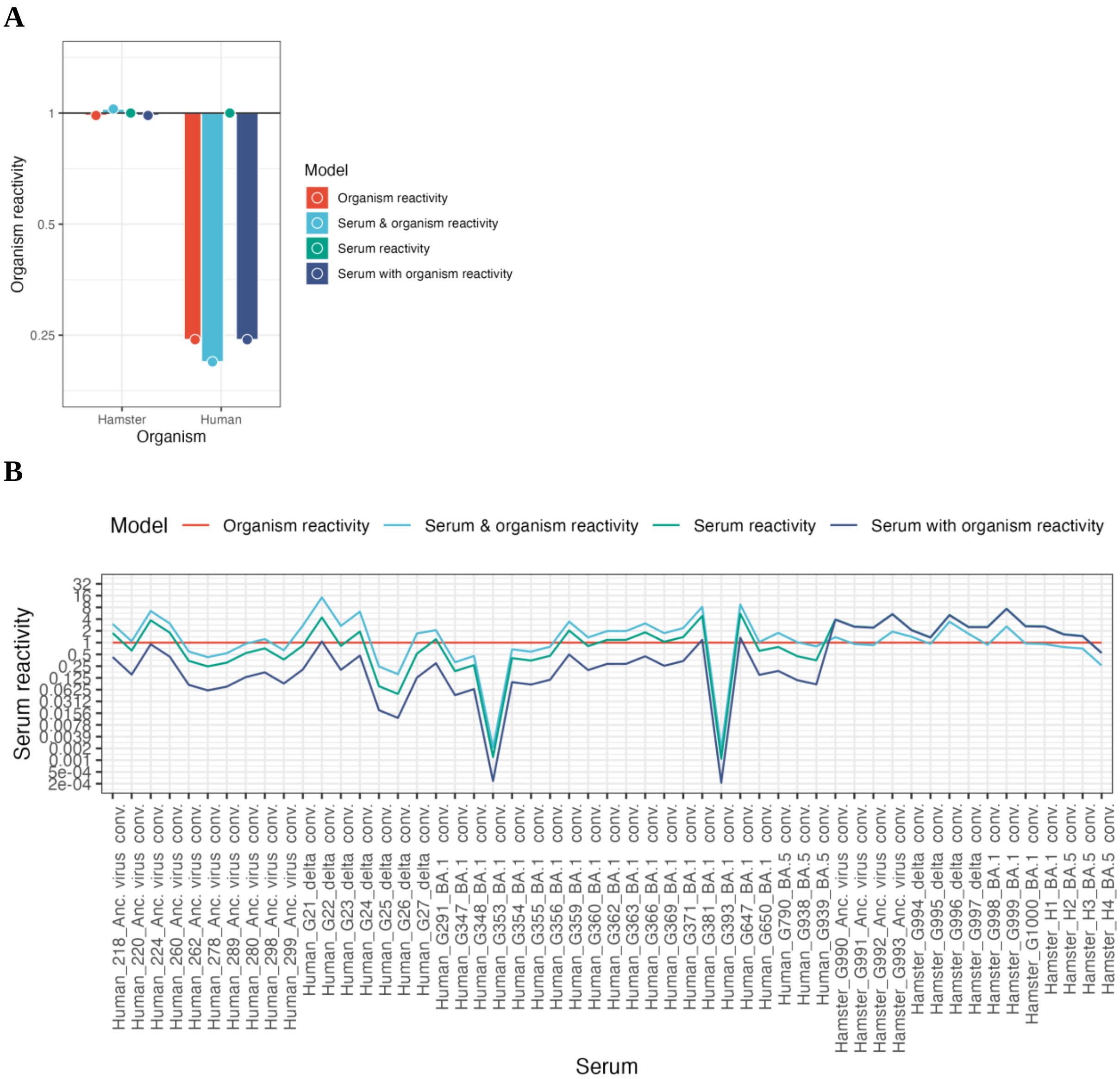


**Fig. S5.**

**Mean organism and serum reactivity effects.** The mean of the sampled posterior distributions for **A)** organism reactivities and **B)** serum reactivities are shown. The red color shows mean values when only organism reactivities were estimated, the green when only serum reactivities were estimated, the light blue when both organism and serum reactivities were estimated, and the dark blue shows the estimated serum reactivities (**B** green) with added estimated organism reactivities (**A** red). The reactivity corresponds to a fold change of titers against all antigen variants.

**Table S1.**
Sample characteristics.

| Species | Exposed Virus Variant | Number | Time since positive PCR/infection† |
|---|---|---|---|
| Human | Ancestral | 10* | 2 – 6 weeks ‡ |
| Human | Delta | 7 | 5 weeks (4 - 18) |
| Human | BA.1 Omicron | 17 | 2 weeks (1.7 - 2.4) |
| Human | BA.5 Omicron | 3 | 3 weeks (2.7 - 3) |
| Hamster | Ancestral | 4 | 26 days |
| Hamster | Delta | 4 | 26 days |
| Hamster | BA.1 Omicron | 4 | 26 days |
| Hamster | BA.5 Omicron | 3 | 21 days |

*5 of the 10 human ancestral variant exposed sera and 2 of the 17 human BA.1 exposed sera were not tested against XBB.1.5.1 due to limited sample volume; †Intervals between positive PCR and blood collection for the different human serum groups were expressed by median and interquartile range (IQR) in weeks when exact intervals were known. Interval between infection and sample collection for the hamster sera are given in days; ‡For the human ancestral convalescent samples, intervals were given as period in weeks, as exact intervals were not known.

**Table S2.**
Virus variants used.

| **Isolate ID*** | **Variant** | **Pango lineage**† | **WHO nomenclature** | **GISAID ID** |
|---|---|---|---|---|
| B86.2 | D614G (ancestral) | B.1.177 | D614G | EPI_ISL_3305837 |
| C63.1 | Alpha | B.1.1.7 | Alpha | EPI_ISL_3277382 |
| C79.2 | Alpha-E484K | B.1.1.7 (E484K) | Alpha (E484K) | EPI_ISL_3277383 |
| C24.1 | Beta | B.1.351 | Beta | EPI_ISL_17528983 |
| D94 | Gamma | P.1.1 | Gamma | EPI_ISL_2095177 |
| D27 | Delta | B.1.617.2 | Delta | EPI_ISL_2290769 |
| E16.1 | BA.1 Omicron | BA.1 | Omicron BA.1 | EPI_ISL_17528984 |
| E65.1 | BA.2 Omicron | BA.2 | Omicron BA.2 | EPI_ISL_12486408 |
| E73.1 | BA.5 Omicron | BA.5.3.2 | Omicron BA.5 | EPI_ISL_13666092 |
| G37.2 | XBB.1.5 Omicron | XBB.1.5.1 | XBB.1.5.1 | EPI_ISL_17077093 |

*Internal name of isolate; †determined using UShER (https://genome.ucsc.edu/cgi-bin/hgPhyloPlace) on 24.03.2023
